# Spectrum of genes for inherited hearing loss in the Israeli Jewish population, including the novel human deafness gene *ATOH1*

**DOI:** 10.1101/2020.06.11.144790

**Authors:** Zippora Brownstein, Suleyman Gulsuner, Tom Walsh, Fábio Tadeu Arrojo Martins, Shahar Taiber, Ofer Isakov, Ming K. Lee, Mor Bordeynik-Cohen, Maria Birkan, Weise Chang, Silvia Casadei, Nada Danial-Farran, Amal Abu-Rayyan, Ryan Carlson, Lara Kamal, Ásgeir Örn Arnþórsson, Meirav Sokolov, Dror Gilony, Noga Lipschitz, Moshe Frydman, Bella Davidov, Michal Macarov, Michal Sagi, Chana Vinkler, Hana Poran, Reuven Sharony, Nadra Samara, Na’ama Zvi, Hagit Baris-Feldman, Amihood Singer, Ophir Handzel, Ronna Hertzano, Doaa Ali-Naffaa, Noa Ruhrman-Shahar, Ory Madgar, Efrat Sofrin, Amir Peleg, Morad Khayat, Mordechai Shohat, Lina Basel-Salmon, Elon Pras, Dorit Lev, Michael Wolf, Eirikur Steingrimsson, Noam Shomron, Matthew W. Kelley, Moien Kanaan, Stavit Allon-Shalev, Mary-Claire King, Karen B. Avraham

**Affiliations:** Department of Human Molecular Genetics & Biochemistry, Sackler Faculty of Medicine and Sagol School of Neuroscience, Tel Aviv University, Tel Aviv, Israel; Departments of Genome Sciences and Medicine, University of Washington, Seattle, WA, USA; Department of Cell and Developmental Biology, Sackler Faculty of Medicine, Tel Aviv University, Tel Aviv, Israel; Raphael Recanati Genetic Institute, Rabin Medical Center–Beilinson Hospital, Tel Aviv University Felsenstein Medical Research Center, Petach Tikva, Israel; Laboratory of Cochlear Development, National Institute on Deafness and Other Communications Disorders, NIH, Bethesda, MD, USA; Genetics Institute, Ha’Emek Medical Center, Afula, Israel; Rappaport Faculty of Medicine, Technion, Haifa, Israel; Department of Biological Sciences, Bethlehem University, Bethlehem, Palestine; Department of Biochemistry and Molecular Biology, BioMedical Center, Faculty of Medicine, University of Iceland, Reykjavik, Iceland; Department of Otolaryngology - Head and Neck Surgery, Schneider Children’s Medical Center, Petach Tikva, Israel; Department of Otolaryngology - Head and Neck Surgery, Sheba Medical Center, Tel Hashomer, Israel; Institute of Human Genetics, Sheba Medical Center, Tel Hashomer, Israel; Department of Human Genetics and Metabolic Diseases, Hadassah-Hebrew University Medical Center, Jerusalem, Israel; Institute of Medical Genetics, Wolfson Medical Center, Holon, Israel; Genetics Institute, Meir Medical Center, Kfar Saba and Sackler Faculty of Medicine, Tel Aviv University, Tel Aviv, Israel; Ziv Medical Center, Zefat, Israel; Genetics Institute, Tel-Aviv Sourasky Medical Center, Tel Aviv, Israel; Community Genetics Department, Public Health Services, Ministry of Health, Ramat Gan, Israel; Department of Otolaryngology Head and Neck Surgery and Maxillofacial Surgery, Tel-Aviv Sourasky Medical Center, Tel Aviv, Israel; Department of Otorhinolaryngology Head and Neck Surgery, University of Maryland School of Medicine, Baltimore, MD, USA; Human Genetics Institute, Lady Davis Carmel Medical Center, Haifa, Israel; Sheba Cancer Research Center, Sheba Medical Center, Tel Hashomer, Israel; Institute of Medical Genetics, Maccabi HMO, Rehovot, Israel

**Keywords:** Next-generation sequencing, massively parallel sequencing, diagnostics, hearing, deafness, gene panel, genomics

## Abstract

Mutations in more than 150 genes are responsible for inherited hearing loss, with thousands of different, severe causal alleles that vary among populations. The Israeli Jewish population includes communities of diverse geographic origins, revealing a wide range of deafness-associated variants and enabling clinical characterization of the associated phenotypes. Our goal was to identify the genetic causes of inherited hearing loss in this population, and to determine relationships among genotype, phenotype, and ethnicity. Genomic DNA samples from informative relatives of 88 multiplex families, all of self-identified Jewish ancestry, with either non-syndromic or syndromic hearing loss, were sequenced for known and candidate deafness genes using the HEar-Seq gene panel. The genetic causes of hearing loss were identified for 60% of the families. One gene was encountered for the first time in human hearing loss: *ATOH1* (Atonal), a basic helix-loop-helix transcription factor responsible for autosomal dominant progressive hearing loss in a five-generation family. Our results demonstrate that genomic sequencing with a gene panel dedicated to hearing loss is effective for genetic diagnoses in a diverse population. Comprehensive sequencing enables well-informed genetic counseling and clinical management by medical geneticists, otolaryngologists, audiologists, and speech therapists and can be integrated into newborn screening for deafness.

## INTRODUCTION

Hearing loss is a leading cause of disability worldwide, with an estimated 466 million people suffering from a loss of greater than 40dB.^1–3^ Hearing loss can have dramatic effects on communication, levels of education, and psychosocial development; it is responsible for a subsequent decline in quality of life, particularly in an increasingly older population.^4,5^ Determining the causes of hearing loss is crucial for clinical management, genetic counseling, and potential prevention. More than 150 genes harbor variants causing non-syndromic hearing loss,^6–8^ and hundreds of genetic syndromes include hearing impairment.^9^ Virtually every population harbors deafness-causing alleles with worldwide distribution and others specific to the local population.^10^

The Jewish population of modern Israel is made up of communities that differ with respect to geographic origin, spoken language, and traditions. Ashkenazi Jews from Europe and North America, Sephardic Jews from North Africa (Morocco, Algeria, Libya, and Tunisia) and southern Europe (Italy, Greece, and Turkey), and Mizrahi Jews from the Middle East (Iran, Iraq, Syria, Yemen, and Egypt) all derive from the Jews who lived in the Middle East 4000 years ago, dispersing with the Babylonian exile in 586 BCE. ^11,12^ After the formation of the state of Israel in 1948, Jews from all these regions immigrated to the country. Today, roughly half of Jewish people live in Israel, yielding an Israeli Jewish population that is approximately 47% Ashkenazi, 30% Sephardi, and 23% Mizrahi.^13,14^ In a study conducted in Israel a few years after its founding, high rates of consanguineous marriage were observed, with the lowest rate (2.5%) among Ashkenazi Jews, and higher rates among non-Ashkenazi Jews, with the highest prevalence (29%) among Jews from Iraq.^15^ During the intervening years, inter-community marriages have become frequent, and consanguineous marriages are much less common.^16^

Centuries of endogamy within each of these communities led to high frequencies of recessive genetic traits, many due to community-specific founder mutations.^13,17,18^ The idea of that each Jewish ancestry has its own genetic blueprint is supported by studies revealing ancestry-specific polymorphisms and haplotypes. Among these are mutations for at least 40 diseases that are specific to individual Jewish communities.^14^ Population-specific mutations include some responsible for hearing loss, such as *GJB2* c.167delT and *TMC1* p.Ser647Phe, while other deafness-causing mutations, such as *GJB2* c.35delG, are common in all Jewish ethnicities and elsewhere^17,1818,16^.

*GJB2* variants are the most prevalent cause of hereditary hearing loss worldwide and are responsible for ~30% of deafness in Jewish families.^17,19,20^ Hence in Israel, routine genetic testing has been for the two most common pathogenic variants, *GJB2* c.35delG and *GJB2* c.167delT. For hearing loss not explained by these alleles, high-throughput sequencing using hearing-loss-dedicated gene panels offers the opportunity to identify other disease-causing variants in hundreds of genes.^21,22,18,23^

The goal of this project was to identify the genetic causes of hearing loss in Israeli Jewish families with more than one affected relative (i.e. multiplex families) and to determine the number of genes responsible for hearing loss in the Israeli Jewish population as a whole. The long-term goal is to apply these results to development of guidelines for the molecular diagnosis of deafness in this population.

## MATERIALS AND METHODS

### Participants

Probands with hearing loss and their relatives were recruited from medical genetics clinics throughout Israel. All probands had a positive family history of hearing loss. Families were asked about their medical history, including family history of relevant symptoms, consanguinity, degree and symmetry of hearing loss, age of onset, use of hearing aids or cochlear implant, tinnitus, exposure to ototoxic drugs or noise, pathological conditions of the inner ear, and vestibular function. Hearing loss could be non-syndromic or syndromic, stable or progressive, and pre-lingual or post-lingual in onset. Probands or their parents gave written informed consent and provided a blood sample for DNA. *GJB2* and *GJB6* were evaluated by Sanger sequencing, and probands with hearing loss due to *GJB2* were so advised and not sequenced with the gene panel.^17^ After these steps, 188 individuals from 88 multiplex families were evaluated with the HEar-Seq gene panels. Hearing controls from each Israeli Jewish ethnic group were identified from healthy, hearing individuals undergoing genetic screening at the Rabin and Sheba Medical Centers and from the National Laboratory for the Genetics of Israeli Populations (https://en-med.tau.ac.il/nlgip). An independent series of 105 individuals with hearing loss, treated at genetics clinics and not related to the 88 multiplex families, were included in order to estimate allele frequencies among Israeli Jewish deaf individuals.

### Experimental procedures

Genomic analysis was carried out with the Hear-Seq gene panels, a custom design by the authors with version ranging from 178 to 372 genes ^18,23,24^ (Table S1). The BED file of Hear-Seq capture probe locations is freely available from the authors, and capture probes can be ordered directly from the manufacturer (Agilent Technologies, Santa Clara, CA, USA) with permission from the authors. Details of panel design and manufacture are included in Supporting Information.

Details of genomic analysis with the panels, of follow-up sequencing, and of the accompanying bioinformatics pipeline are also included in Supporting Information. Novel mutations of uncertain consequence in *ATOH1* and *MITF* were evaluated by protein biochemistry and cell biology. Details of these methods are also included in Supporting Information.

## RESULTS

### Genetic diagnoses of hearing loss from gene panel sequencing

Genetic causes of hearing loss were identified for 60% (53/88) of the families evaluated by the HEar-Seq gene panels. These genetic diagnoses involved 57 different causal alleles in 27 different genes (Table S2, Figure S1). Most of the responsible alleles (32 of 57, or 56%) had not been previously reported from any population (Table 1). Of the novel variants, 50% were missense, 31% frameshifts, 9% nonsense, 9% copy number variants (CNVs), and 3% silent mutations that altered splicing. These diagnoses expanded the total number of genes known to be responsible for inherited loss in the Israeli Jewish population from seven^17^ to 32 (Table S3).

**TABLE 1.** Novel variants discovered by the HEar-Seq gene panel in Israeli Jewish families with hearing loss

| Gene | Family | Inheritance* | Ethnicity** | Genomic coordinates (hg19) | cDNA | Variant type | Effect | Allele freq controls of same ethnicity* | Allele freq in unrelated deaf^^ | ClinVar ID | ACMG classification (criteria)*** |
| --- | --- | --- | --- | --- | --- | --- | --- | --- | --- | --- | --- |
| *ATOH1*<br>NM_005172.1 | HL263 | AD | M | chr4:94751102 | c.1030delC | Indel | His344fs17Ter | 0 | 0 | 813817 | P (PM2, PM4, PP1_S, PP3, PP4) |
| *ATP2B2*<br>NM_001001331.2 | DF328 | AD | A | chr3:10420936 | c.1033C>T | Nonsense | Gln345Ter | 0 | 0 | 1337668 | P (PM2, PM4, PP1_S, PP3, PP4) |
| *CABP2*<br>NM_001318496.1 | DF326 | AR | Mixed | chr11:67289435 | c.250G>A | Missense | Glu84Lys | 0 | 0 | 1337669 | P (PM2, PM3_S, PP1_M, PP3, PP4) |
| *CEACAM16*<br>NM_001039213.3 | DF301 | AR | M | chr19:45208901<br>Founder allele | c.703C>T | Missense | Arg235Cys | 0 | 0.008 | 236048 | P (PS4, PM2, PP1_S, PP3, PP4) |
| *CLPP*<br>NM_006012.2 | DF313 | AR | S | chr19:6361758 | c.173T>G (mat) | Missense | Leu58Arg | 0.004 | 0 | 500291 | 1) LP (PM2, PM3_P, PP1_M, PP3, PP4) |
|  |  |  | A | chr19:6361914 | c.233G>C (pat) | Missense | Arg78Pro | 0 | 0 | 813818 | 2) LP (PM2, PM3_P, PP1_M, PP3, PP4) |
| *COCH*<br>NM_004086.2 | HL1103 | AD | A | chr14:31355200 | c.1159C>T | Missense | Leu387Phe | 0 | 0 | 236036 | LP (PM2, PP1_M, PP3, PP4) |
| *COL11A2*<br>NM_080680.2 | HL1140 | AR | S | chr6:33152074 | c.967insC | Indel | Thr323fs17Ter | 0 | 0 | 813820 | 1) P (PVS1, PM2, PM3_P, PP1_M) 2) P (PM2, PM3_S, PP1_M, PP3, PP4) |
|  |  |  | M | chr6:33138676 | c.3385G >A | Missense | Gly1129Arg | 0 | 0 | 813821 |  |
| *EYA4*<br>NM_172105.3 | HL21 | AD | Mixed | chr6:133783474 | c.441delC | Indel | Tyr148fs49Ter | 0 | 0 | 236032 | P (PVS1, PM2, PM4, PP1, PP3, PP4) |
| *EYA4*<br>NM_172105.3 | DF312 | AD | M | chr6:133844297 - 133844299 | c.1720_1722 delTACinsAAA | 2 bp changes in same codon | Tyr574Lys | 0 | 0 | 1164280 | LP (PM2, PM2, PP1, PP3, PP4) |
| *GATA3*<br>NM_002051.2 | HL738 | AD | A | chr10:8100707 | c.681ins35 | Indel | Glu228fs37Ter | 0 | 0 | 236031 | P (PVS1, PM2, PP1_M, PP3, PP4) |
| *GATA3*<br>NM_002051.2 | HL769 | AD | Mixed | chr10:8106009 | c.829G>A | Missense | Asp277Asn | 0 | 0 | 813823 | LP (PM2, PP1_S, PP3) |
| *MITF*<br>NM_198159.2 | DF311 | AD | S | chr3:70001035 | c.935T>C | Missense | Leu312Pro | 0 | 0 | 547531 | P (PS4_P, PM2, PP1, PP3, PP4) |
| *MITF*<br>NM_198159.2 | DF186 | AD | S | chr3:70014025 | c.1190delG | Indel | Gly397fs15Ter | 0 | 0 | 236050 | P (PVS1, PM2, PP1, PP3, PP4) |
| *MITF*<br>NM_198159.2 | DF219 | AD | A | chr3:70005649 | c.981InsC | Indel | Leu327fs9Ter | 0 | 0 | 813825 | P (PVS1, PM2, PM4, PP1, PP3, PP4) |
| *MYO6*<br>NM_004999.3 | DF305 | AD | A | chr6:76538307 | c.238C>T | Nonsense | Arg80Ter | 0 | 0 | 178957 | P (PVS1, PM2, PP1, PP3, PP4) |
| *MYO6*<br>NM_004999.3 | HL1133 | AD | S | chr6:76568683 | c.1452insT | Nonsense | Asn485Ter | 0 | 0 | 1337671 | P (PVS1, PM2, PM4, PP1_M, PP3, PP4) |
| *MYO6*<br>NM_004999.3 | HL1274 | AD | A | chr6:76568710 | c.1473del3insC | Indel | skip exon 14,<br>Glu461fs13Ter | 0 | 0 | 236034 | P (PVS1, PM2, PM4, PP1, PP3) |
| *MYO6*<br>NM_004999.3 | HL158 | AD | A | chr6:76624636 | c.3765delC | Indel | Cys1256fs28Ter | 0 | 0 | 813826 | P (PVS1, PM2, PM4, PP1, PP3, PP4) |
| *MYO15A*<br>NM_016239.3 | HL72 | AR | M | chr17:18054799 | c.7751_8224del3446ins23 | CNV | Gln2583fs19Ter | 0 | 0.003 | 236039 | P (PVS1, PM2, PM4, PP1, PP3, PP4) |
| *MYO15A*<br>NM_016239.3 | DF327<br>DF317 | AR | A | chr17:18069748<br>Founder allele | c.9861C>T | Silent | Gly3287Gly | 0 | 0.003 | 228276/45777 | P (PM2_P, PM3_S, PP1_S, PP3) |
| *PCDH15*<br>NM_001142769.1 | HL1134 | AR | M | chr10:56287598 | c.146T>C | Missense | Val49Ala | 0 | 0.003 | 450626 | LP (PM2, PM3_P, PP1, PP3, PP4) |
| *SLC26A4*<br>NM_000441.1 | HL1132<br>HL1327 | AR | A | chr7:107312627 | c.349C>T | Missense | Leu117Phe | 0.007 | 0.013 | 43555 | P (PS4, PM1, PM3, PM3_P, PP1_S, PP3, PP4) |
| *SOX10*<br>NM_006941.3 | HL971 | AD | A | chr22: 38379660 | c.125_132del8 | Indel | Leu42fs21Ter |  | 0 | 813829 | P (PVS1, PM2, PM4, PP3, PP4) |
| *STRC*<br>NM_153700.2 | HL927 | AR | Mixed | chr15:43,892,353 - 43,910,998 (min) | del entire gene | CNV | - |  | 0.003 | 1164296 | P (PVS1, PM1, PM3, PM4, PP1, PP3, PP4) |
| *TECTA*<br>NM_005422.2 | DF303 | AR | A | chr11:120979969 | c.248C>T | Missense | Thr83Met |  | 0.003 | 813831 | LP (PM2, PM3_P, PP1_M, PP3, PP4) |
| *TECTA*<br>NM_005422.2 | HL277 | AD | S | chr11:121000866 | c.2887G>A | Missense | Ala963Thr | 0 | 0 | 236059 | LP (PM2, PP1_M, PP3, PP4) |
| *TECTA*<br>NM_005422.2 | DF183 | AD | A | chr11:121058558 | c.6017A>G | Missense | Asp2006Gly | 0 | 0 | 236033 | LP (PM2, PM5, PP1, PP3, PP4) |
| *TJP2*<br>NM_001170414.2 | DF180 | AD | S | chr9:71704982 - 71840362 dup | dup exons 1-6 | CNV | - | 0 | 0 | 236035 | P (PS4_P, PM1, PM2, PM4, PP1_S, PP3, PP4) |
| *TMC1*<br>NM_138691.2 | DF193 | AR | Mixed | chr9:75263573 | c.15insA | Indel | Val6fs12Ter | 0 | 0.003 | 236041 | P (PVS1, PM2, PM4, PP1, PP3, PP4) |
| *TMC1*<br>NM_138691.2 | HL1159 | AR | A | chr9:75369733 | c.674C>T | Missense | Pro225Leu | 0 | 0 | 424807 | P (PM2, PM3_S, PP1_M, PP3, PP4) |

\*Recessive, R; Dominant, D
\*\* A, Ashkenazi; M, Mizrahi; S, Sephardi
\*\*\*ACMG classification and criteria legend: P, pathogenic; LP, likely pathogenic; PVS, very strong pathogenicity evidence; PS, strong pathogenicity evidence; PM, moderate pathogenicity evidence; PP, supporting pathogenicity evidence; \_S, strong; \_M, moderate; \_P, supporting ACMG classification and criteria<sup>1,2</sup>
^Number of ethnicity-matched hearing controls ranges from 163-500 individuals (326-1000 chromosomes)
^^Number of unrelated deaf is 105 individuals (210 chromosomes)

### *ATOH1*, a new gene for human hearing loss

Panel sequencing revealed involvement of a new gene for human hearing loss. *ATOH1* (Atonal) encodes a basic helix-loop-helix transcription factor that is essential for neuronal development in the cerebellum.^25^ Heterozygosity for any of several different variants of *Atoh1* leads to hearing deficits in mice, some with syndromic features.^26,27^ *ATOH1* was included on the panel because mutations in the mouse ortholog lead to hearing loss. In Iraqi Jewish kindred HL263, *ATOH* c.1030delC co-segregated over five generations with progressive non-syndromic hearing loss, with onset at birth or early childhood (Figure 1A,B). Based on whole exome sequencing, no other potentially damaging variant in any gene co-segregated with hearing loss in this family. *ATOH* c.1030delC causes a frameshift that alters the last ten residues of the normally 354-amino acid protein and adds six residues to its length before a stop.

**Figure 1.**
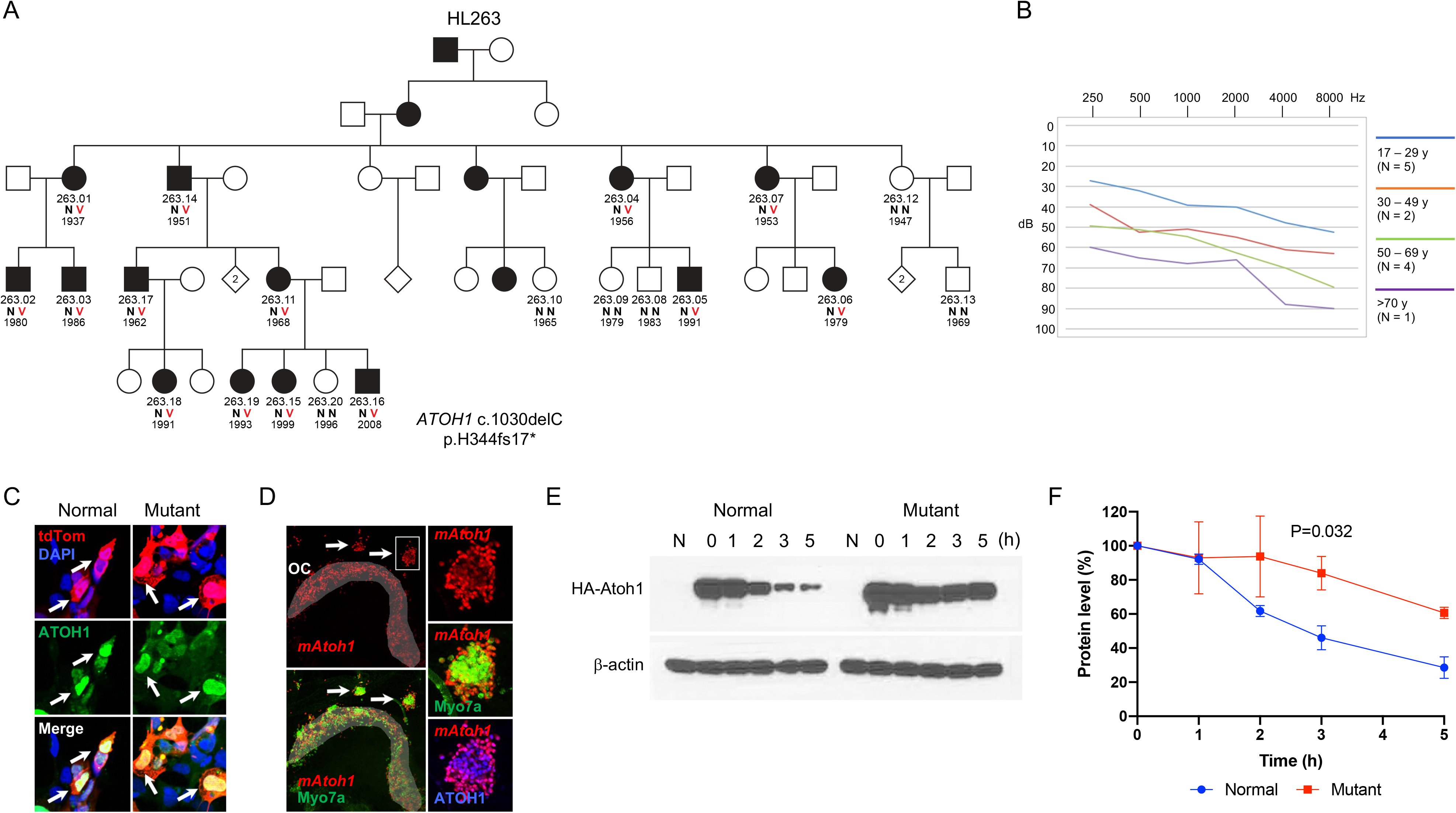
*ATOH1 c.1030delC* and age-related hearing loss. **A**, Family HL263 with progressive sensorineural hearing loss in five generations. Filled symbols represent individuals with hearing loss. V represents the variant allele and N the normal allele. The number under each individual is the birth year. **B**, Average hearing thresholds of family members of various ages heterozygous for the mutation. **C**, HEK293 cells transfected with expression constructs for *ATOH1*^*WT*^ or *ATOH1*^*c.1030delC*^. Anti-ATOH1 labeling in green indicates nuclear localization of both wild type and mutant ATOH1 (arrows). **D**, left column: Cochlear explant from an *Atoh1*^−/−^ mouse established at E14 and transfected with an *ATOH1*^*c.1030delC*^ expression construct. Transfected cells (red) are present in a region that is consistent with the location of the organ of Corti in a wildtype cochlea (shaded region, OC), as well as in several patches that are located outside of the organ of Corti (arrows). Counter-staining for the hair cell marker Myo7a indicates that many transfected cells have developed as hair cells. Right column: High magnification images of the boxed region demonstrating expression of Myo7a (green) and ATOH1 (blue) in a patch of transfected cells. **E**, Western blot of ATOH1 protein extracted from HEK293T cells transfected with wild type or mutant ATOH1, after 1-5 hours treatment with 1mM cycloheximide. **F**, Quantification of the results of part C. Statistical test was repeated measures ANOVA with post-hoc Holm-sidak correction for multiple comparisons.

To examine the effects of this mutation, expression constructs for wild type and mutant *ATOH1* were transfected into HEK293 cells. Antibody localization indicated nuclear translocation of both wild-type and mutant *ATOH1* (Figure 1C). Next, to determine the effects of the mutation on protein function, cochlear explants were established at E14 or E15 from *Atoh1*^−/−^ animals and transfected with either the *ATOH1*^*WT*^ or *ATOH1*^*c.1030delC*^ expression construct. (*Atoh1*^−/−^ cochleae were used to ensure no potential effects from the endogenous gene.) Results indicated induction of Myo7A^+^ hair cells in response to expression of either construct (Figure 1D). However, western analysis revealed a significantly slower rate of degradation for mutant ATOH1 protein compared to wild-type ATOH1 protein (P=0.032, multiple t-test with Holm-Sidak correction, two biological replicates) (Figure 1E, F).

### Complex relationships of genotypes to phenotypes

For genes responsible for syndromic hearing loss, different variants in the same gene revealed new relationships of genotypes to phenotypes. Three families with mutations in *MITF* illustrate these complexities (Figure 2). Damaging variants of *MITF* can cause autosomal dominant Waardenburg type 2A and Tietz albinism/deafness syndromes, both of which are highly heterogeneous clinically (Figure 2A, B). In family DF311, three relatives heterozygous for *MITF* c.935T>C, p.(Leu312Pro) had severe to profound sensorineural hearing loss with congenital albinism. (Hearing loss of DF311.02 is due to a mutation in *CDH23*.) Leu312 is a completely conserved residue in the middle of the MITF basic helix-loop-helix (bHLH) domain. A proline at this position would likely break the helix and preclude proper DNA binding and possibly preclude dimerization as well. In family DF219, three relatives heterozygous for *MITF* c.981insC, p.(Leu327fs9Ter) demonstrated the same hearing loss and albinism. A frameshift at residue 327 would lead to truncation in the middle of the bHLH domain and loss of normal protein function. In contrast, in family DF186, three relatives heterozygous for *MITF* c.1190delG, p.(Gly397fs15Ter) also demonstrated congenital sensorineural hearing loss but no pigmentation signs other than hair whitening of the mother in her twenties. Truncation due to frameshift at residue 397 is distal to the bHLH domain, so its consequences to protein function were unknown. A transactivation assay of the protein encoded by *MITF* c.1190delG, p.(Gly397fs15Ter) indicated that the transcriptional potential of the mutant protein is greatly impaired compared to that of wild-type MITF (Figure 2C, D).

**Figure 2.**
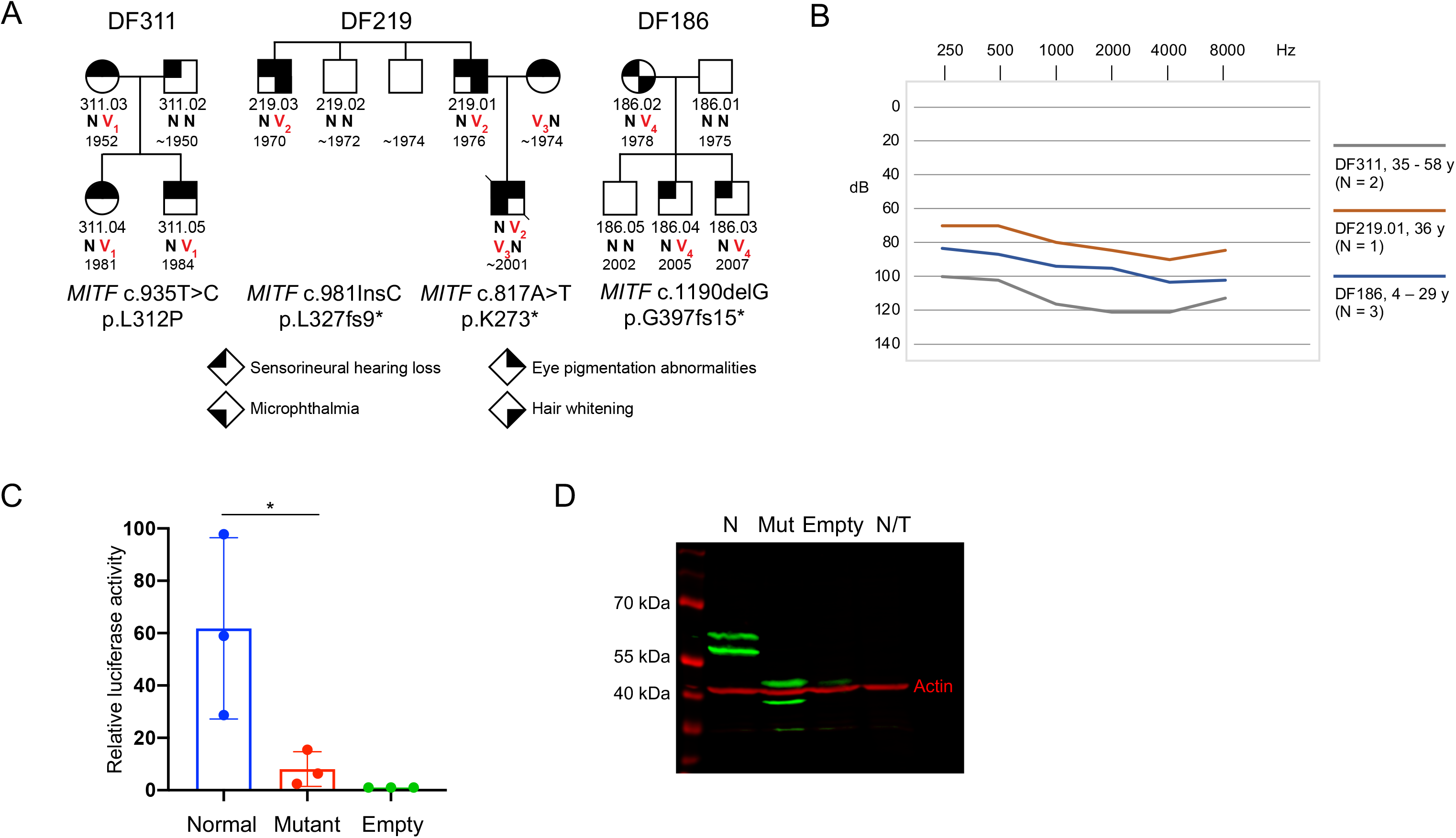
*MITF1* variants associated with hearing loss and Waardenburg Syndrome Type 2A / Tietz Syndrome in three families. **A**, Pedigrees of families DF311, DF186, and DF219, indicating variation in syndromic features. **B**, Hearing thresholds by age, reflecting severe to profound hearing loss in all affected individuals. **C**, Transactivation assay using a tyrosinase promoter and luciferase reporter revealing that the transcriptional potential of protein encoded by *MITF* c.1190delG is greatly impaired compared to the wild-type protein. Statistical test was one-tailed student’s t-test. **D**, Western blot analysis indicates similar levels of wild type and mutant MITF protein.

*GATA3* variants present an analogous story. *GATA3* is responsible for an autosomal dominant syndrome including hypoparathyroidism, sensorineural deafness, and renal dysplasia (HDR), for which different alleles are associated with a wide spectrum of phenotypes.^28^ Two families in our series reflect this heterogeneity (Figure 3). In family HL738, the proband (DF738.01), heterozygous for *GATA3* c.681ins35, p.(Glu228fs37Ter), and his mother both demonstrated congenital severe-to-profound hearing loss and kidney dysplasia. (Medical records for the sister and niece of the proband were not available.) In contrast, in family HL769, all relatives heterozygous for *GATA3* c.829G>A, p.(Asp277Asn), demonstrated severe-to-profound hearing loss, but to date no renal or parathyroid problems.

**Figure 3.**
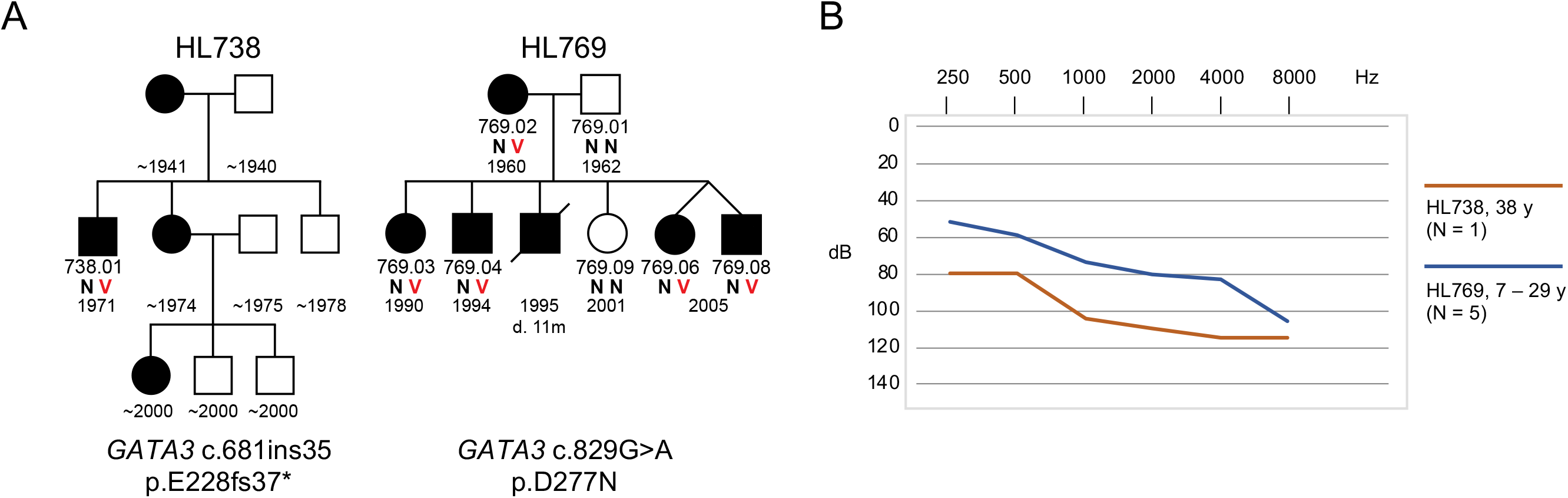
*GATA3* variants associated with nonsyndromic and syndromic hearing loss in two families. A, Pedigrees of families HL738 and HL769. B, Hearing thresholds by age, reflecting severe to profound hearing loss in all affected individuals.

*CLPP* is associated with Perrault syndrome, characterized by sensorineural hearing loss (SNHL) and infertility in both females and males.^29^ In family DF313, affected siblings were both compound heterozygous for novel variants *CLPP* c.173T>G, p.(Leu58Arg) and c.233G>C, p.(Arg78Pro) (Figure S1). These siblings are young, age 3y, and thus far have profound hearing loss and auditory neuropathy but no other symptoms. It is not clear whether they are pre-symptomatic for infertility or if this genotype leads to nonsyndromic hearing loss.

*SOX10* is responsible for autosomal dominant Waardenburg syndrome types 2E^30^ and 4C,^31^ and for Kallmann syndrome.^32^ The adult proband of family HL971 and his mother, both with congenital profound hearing loss, are heterozygous for novel variant *SOX10* c.125_132del8, p.(Leu42fs21Ter) (Figure S1). Neither the proband nor his mother initially reported any symptoms other than deafness. Upon detecting this *SOX1* variant and asking the proband about his sense of smell, we discovered that both he and his deaf mother have anosmia, characteristic of Kallmann syndrome.

*USH2A* is responsible for Usher syndrome type 2A but signs of retinitis pigmentosa (RP) depend on genotype and on age. In family HL149, the proband, age 3y is compound heterozygous for *USH2A* c.3368A>G, p.(Tyr1123Cys)^33^ and *USH2A* c.240_241insGTAC, a known pathogenic variant associated with SHL^34^ (Table S2). His mother, in her early thirties, is homozygous for USH2A c.240_241insGTAC, with sloping mild-to-severe hearing loss and mild RP. The proband also has a sloping mild-to moderate hearing loss, but as yet no signs of RP.

*TECTA* can be responsible for autosomal dominant or autosomal recessive hearing loss. Among families with dominant hearing loss, missense mutations in the TECTA zonadhesion domain (amino acid residues 260-1694) are associated with high-frequency hearing loss, while missense mutations in the TECTA zona pellicuda domain (residues 1795-2059) are associated with mid-frequency hearing impairment^35–37^. This correspondence obtains for family HL277 (p.Ala963Thr) and family DF193 (p.Asp2006Gly) (Figure S1).

### Contributions of founder alleles

Founder alleles of ancestral Jewish communities continue to contribute to hearing loss in Israeli Jewish families. Table S3 lists the 75 variants in 32 genes of the families in the study, distributed among different communities. Some variants were private to one family, whereas others were more common but limited to one community, reflecting a founder effect (Table S4). Among founder mutations, the principal contribution to hearing loss was of course from *GJB2* alleles, but founder mutations in other genes also contributed. For example, *CEACAM16* c.703C>T, p.(Arg235Cys) was responsible for recessive hearing loss in family DF301, of Jewish Iranian ancestry (Figure S1). This allele was subsequently identified in children with hearing loss from other families of Jewish Iranian ancestry evaluated in clinics. Another example is *OTOF* c.5193-1G>A, responsible for recessive hearing loss with auditory neuropathy in family HL1015, of Syrian Jewish ancestry (Figure S1). This allele was heterozygous in three of 184 hearing controls of Syrian Jewish ancestry (allele frequency 0.008), but was absent from hearing controls of all other Jewish ethnicities and absent from gnomAD. It is likely a founder mutation among Syrian Jews.

A quite common founder allele in the Ashkenazi community is *SLC26A4* c.349C>T, p.(Leu117Phe), with allele frequency 0.005 in the Ashkenazi Jewish population compared to 0.0002 in other gnomAD populations. No homozygotes for this variant have been observed among hearing individuals of any ancestry. This missense occurs in a completely conserved residue in the first transmembrane domain of SLC26A4, but its consequence has been uncertain. Families HL1132 and HL1327 were informative for this variant (Figure S1). Of the five children with hearing loss in these families, four are homozygous for *SLC26A4* p.(Leu117Phe). These deaf children are the first homozygotes reported for this variant. HL1327.09, who has profound hearing loss but is heterozygous for *SLC26A4* p.(Leu117Phe), remains unexplained. She has no other mutation in *SLC26A4*. She may be a phenocopy for inherited hearing loss in this family. Homozygosity for *SLC26A4* p.(Leu117Phe) has also been observed in four other Ashkenazi families during screening for childhood hearing loss by genetics clinics in Israel. We speculate that *SLC26A4* c.349C>T, p.(Leu117Phe), or a non-coding regulatory mutation of *SLC26A4* in tight linkage disequilibrium with it, is pathogenic for nonsyndromic hearing loss.^38–40,41^

Figure 4 presents genotype-phenotype-ancestry associations for all variants encountered in the study.

**Figure 4.**
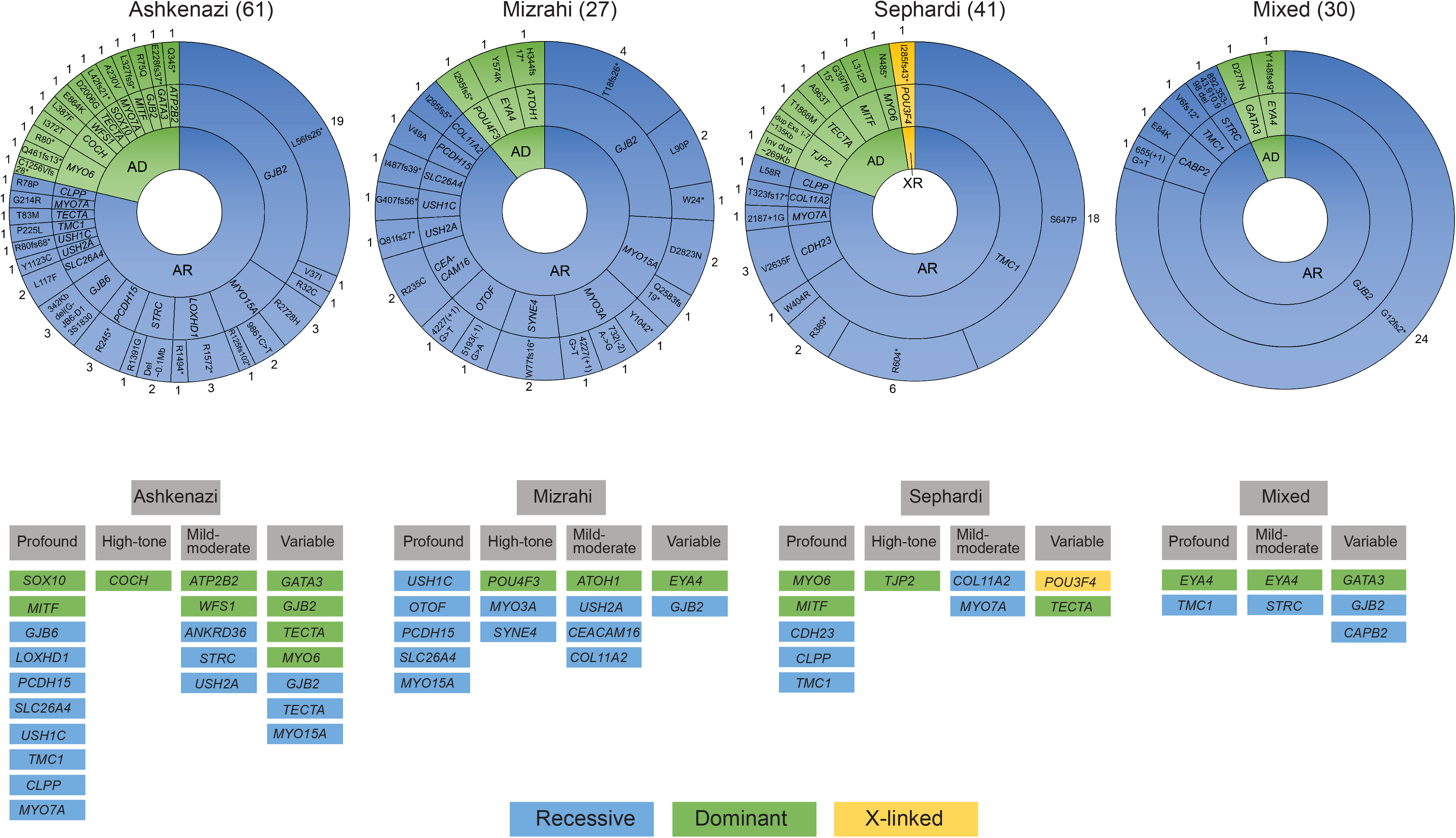
Distribution of hearing loss variants in Ashkenazi, Mizrahi, Sephardi, and mixed Jewish communities. Numbers in parentheses are the number of probands of each ethnicity. Numbers of cases are listed next to each gene.

## DISCUSSION

Genetic diagnosis will play an increasing role in treatment of both congenital and later onset hearing loss. Success of cochlear implant may depend on the genetic cause of the hearing loss.^42^ The clinical application of gene therapy for some forms of hearing loss may prove feasible,^43,44^ but its application depends on correct genetic diagnosis. Gene panel-based sequencing increased the yield of genetic diagnoses from 23%^17^ to 60% of familial hearing loss in the Israeli Jewish population. The analysis revealed 57 different pathogenic variants in 27 genes, with most variants not previously reported, and increased the number of genes known to cause hearing loss in the Jewish population from seven to 32.

Yield from the HEar-Seq gene panel compares favorably with whole exome sequencing (WES).^45^ Costs of WES have decreased in recent years, but far fewer patients can be sequenced simultaneously with high coverage by WES than with a gene panel. Gene panels have proven effective for genetic diagnosis of a tremendous variety of conditions, ranging from inherited predisposition to cancer^46^ to inherited eye disorders.^47^ Gene panel sequencing also minimizes the frequency of incidental findings,^48^ which can introduce legal, ethical, and social dilemmas. Nevertheless, for families not solved by panel sequencing, WES is the next step in searching for a genetic diagnosis.

The discovery that mutation of *ATOH1* can cause human hearing loss adds to understanding the role of this transcription factor in mammalian hearing. ATOH1 is crucial for the development and differentiation of inner-ear hair cells^49^ and is first expressed in the nascent organ of Corti. Loss of *Atoh1* in mice causes hearing impairment, cerebellar and cochlear malformations, and death,^25^ while conditional deletion of *Atoh1* leads to lack of differentiated inner ear hair cells,^50^ and the naturally occurring mutation Atoh1 p.Met200Ile causes hearing loss, progressive cerebellar atrophy, and trembling.^26^ The *ATOH1* mutation of family HL263 yields a protein with an abnormal C-terminus associated with an abnormally slow degradation rate. This is consistent with previous observations that *Atoh1* protein stability is regulated by its interaction with the E3 ubiquitin ligase Huwe1 at phosphorylation sites S328 and S339 (human S331 and S342).^51,52^ In this context, a mouse with mutation at the phosphorylation site Atoh1 S193 was shown to have late-onset deafness.^27^ In wild-type mice, expression of *Atoh1* ceases by the end of the first postnatal week. Induction of *Atoh1* at the neonatal stage causes formation of immature ectopic hair cells, with randomized stereocilia orientation and reduced basolateral measured currents.^53,54^ Atoh1 is also known to positively auto-regulate its own expression.^55^ We showed that the human *ATOH1* mutation increases the stability of the protein through decreased degradation. We hypothesize that persistent untimely expression of ATOH1 may generate immature ectopic hair cells that interfere with development of normal hair cells.^56,57^

Early diagnosis enables caregivers to learn whether children are likely to develop symptoms other than hearing loss and so plan for specialized education and treatment. Among our families, pathogenic variants in *CLPP*, *SOX10*, and *USH2A*, which cause syndromic hearing loss, were detected in children prior to onset of additional symptoms. Similarly, a missense mutation in *GATA3* was present in a family with relatives ages 15-60y with apparently non-syndromic hearing loss. Ages at onset of the syndromic features of *GATA3* are variable, with mean age of onset during childhood or early teenage years, but possible onset of the full spectrum of phenotypes as late as age 50.^58^ It is still possible, therefore, that the affected relatives of this family will develop syndromic features. It is also possible that mild mutations in *GATA3* could cause non-syndromic hearing loss.

In summary, the diversity of the Israeli Jewish population is reflected in the diversity of mutations that have been revealed to be responsible for hearing loss. These variants include mutations private to one family, founder mutations of ancestral communities, and mutations appearing in families of all ancestries. Our results are informative for genetic counselors, medical geneticists, audiologists and otolaryngologists caring for families with inherited hearing loss. Genetic diagnosis can be integrated with history, physical examination, and audiometry to guide management of patients with hearing loss. The results can also assist in developing guidelines for genetic screening of newborns with possible hearing loss, in Israel and elsewhere.

## SUPPORTING INFORMATION

**Table S1**. Genes on HEar-Seq panels

**Table S2**. Variants identified by HEar-Seq in Israeli Jewish families

**Table S3**. All variants leading to hearing loss in Israeli Jewish Families in the study

**Table S4**. Founder mutations in Israeli Jewish families

**Figure S1**. Pedigrees, audiograms and genotypes of families with hearing loss. N = wild type; V = variant.

## ACKNOWLEDGEMENTS

The authors wish to thank all the families for their participation in this study. This research was supported by grants from the National Institutes of Health/National Institute of Deafness and Communication Disorders (NIDCD) R01DC011835 (K.B.A., M.C.K., M.N.K.), the Intramural Program at the NIDCD, DC000059 (M.W.K.), the Israel Precision Medicine Program or the Israel Science Foundation, ISF3499-19 (K.B.A.), the Ernest and Bonnie Beutler Research Program of Excellence in Genomic Medicine (K.B.A.), the Hedrich Charitable Trust (K.B.A.), and travel grants from the University of Washington Virginia Bloedel Hearing Research Institute (K.B.A., R.C.).

## AUTHOR CONTRIBUTIONS

K.B.A. and M.-C.K. had full access to all of the data in the study and take responsibility for data integrity and analytic accuracy. Z.B., S.G, T.W., K.B.A. and M.-C.K. were responsible for the concept and design. Z.B., S.G., T.W., F.T.A.M., S.T., O.I., M.K.L., M.B., W.C., S.C., M.B.-C., N.D.-F., A.A.-R., R.C., L.K., A.O.A., M.S., D.G., N.L., M.F., B.D., M.M., M.S., H.V., H.P., R.S., N.S., N.Z., H.B.-F., A.S., O.H., R.H., D.A.-N., N.R-S., O.M., E.S., A.P., M.K., M.S., L.B.-S., E.P., D.L., N.S., M.W.K., M.-C.K., K.B.A. acquired, analyzed, and/or interpreted the data. Z.B., F.T.A.M., M.B.-C., W.C., E.S., M.-C.K. and K.B.A. drafted the manuscript. S.G., O.I., M.K.L., N.S. and M.-C.K. performed the statistical analysis. All authors approved the final version of the manuscript.

## ETHICS APPROVAL

The Ethics Committee of Tel Aviv University, the Helsinki Committee of the Israel Ministry of Health, and the Human Subjects Division of the University of Washington approved the study. Care and housing of *Atoh1^+/−^* mice was conducted based on the NIH guidelines for animal use (Protocol number 1262).

## DATA AVAILABILITY STATEMENT

Novel variants are available at ClinVar (www.ncbi.nlm.nih.gov/clinvar/).

## Supporting Information

### METHODS

#### Genomics

Genomic DNA was sequenced using HEar-Seq gene panels containing a custom design of 178 to 372 genes developed by the authors and manufactured by Agilent Technologies (Santa Clara, CA, USA) Genes included in each version of Hear-Seq are indicated in Table S1. For each version of the panel, hg19 genomic coordinates were submitted to eArray (Agilent) to design cRNA oligonucleotides to cover exons and flanking introns. The BED file of capture probe locations is freely available upon request, and capture probes can be ordered directly from Agilent Technologies with permission from the authors. HEar-Seq v4 includes 178 protein-coding genes and one miRNA, with a total target size of 808 kb that includes 3134 exons and 25bp of intronic sequence flanking each splice site of each exon. Genes included in all previous versions of Hear-Seq are indicated in Table S1.

For analysis of participant genomic DNA, molecular barcodes were assigned and 96 samples multiplexed and sequenced in a single flow cell of the Illumina HiSeq 2500 with 105bp paired-end reads. The average depth of coverage per sample across the HEar-Seq panel was 638x (range of 161 - 924x), with 99.1% of targeted bases with a minimum coverage of 10x and 96.5% of bases with minimum coverage of 50x. Capture and sequencing using the HEar-Seq panels were performed at the University of Washington or the Technion Genome Center. Genotypes of family members were determined by panel sequencing or Sanger sequencing of the family’s identified variant. Genotypes of controls and unrelated deaf were evaluated by Sanger sequencing or restriction enzyme analysis. For family HL263, whole exome sequencing was carried out to rule out other pathogenic variants.

A bioinformatics pipeline was designed for analysis of gene panel data. Data were collected locally and processed on a Dell PowerEdge R920 server. Samples were processed from real-time base-calls (RTA1.8 software [Bustard]) and converted to qseq.txt files. Following demultiplexing, the reads were aligned to the reference human genome (hg19) using Burrows-Wheeler Aligner (0.7.9a).^1^ PCR duplicates were removed by SAMtools 0.1.19 (http://samtools.sourceforge.net/). Indel realignments and base quality score recalibration were carried out, and genotypes called and filtered, with the Genome Analysis Tool Kit (GATK; v3.0-0; broadinstitute.org/gatk) with recommended parameters.^2^ Large insertions, deletions, and inversions were identified with Pindel 0.2.5^3^ and BreakDancer 1.1.4.^4^ Mis-alignments were corrected by aligning to data from 850 in-house exomes. CNVs were called using CoNIFER,^5^ XHMM^6^ and an in-house CNV detection pipeline.^7^ Variants were annotated using an in-house pipeline with respect to location (exonic, near splice, intronic, UTRs) and predicted function (frameshift, inframe indel, nonsense, missense, silent, and potential splice altering, cryptic splice, and regulatory effect). Allele frequencies were obtained from the gnomAD browser. GERP, PolyPhen2, SIFT, and Mutation Taster scores were included. Potential splice altering and cryptic splice variants were predicted using our splice variant prediction pipeline, based on the NNSPLICE and MaxEnt algorithms.^8,9^ Variants were classified using the criteria of the American College of Medical Genetics (ACMG) criteria^10,11^ and deposited in ClinVar (www.ncbi.nlm.nih.gov/clinvar/).

#### ATOH1 analysis

HEK293 cells were cultured on cover slips coated with gelatin. Wild-type and mutant human *ATOH1* sequences were cloned into a pCdT expression construct that also included an *IRES-tdTomato* sequence. Following confirmation by sequencing, HEK cells were transfected with either expression plasmid using Xfect Transfection Reagent (Clontech). Three days after transfection, cells were fixed for 10 minutes in 4% paraformaldehyde. Immunostaining was carried out with primary antibodies against ATOH1^12^ and MYO7A (Proteus). Antibody labeling was detected using Alexa goat anti-rabbit 633 and Alex goat anti-chicken 488 (Thermofisher) secondary antibodies. Finally, nuclei were labeled using a short incubation with DAPI at 1:10,000 in saline. Expression of tdTOMATO was visualized directly. Labeling was imaged on a Zeiss LSM710.

Localization studies of wildtype and mutant Atoh1 were carried out in cochlear explants from *Atoh1*^+/−^ animals^13^ and obtained from the Jackson Laboratory. Care and housing of animals was conducted based on the NIH guidelines for animal use (Protocol number 1262). *Atoh1*^+/−^ animals were crossed to generate time-pregnant females. At E14 or E15, females were euthanized and embryos were removed. Cochleae from each embryo were dissected into separate dishes containing cold PBS. DNA was extracted from the tail of each embryo using a Maxwell RSC instrument (Promega) and genotyped by PCR. *Atoh1*^−/−^ cochleae identified and then electroporated with either *pCdT ATOH1*^*WT*^-*IRES-tdTom* or *pCdT ATOH1*^*c.1030delC*^-*IRES-tdTom* at 1 ug/ul in water.^14^ The electroporated cochlear explants were incubated for 8-9 days^15^, then fixed with 4% paraformaldehyde for 30 min. Expression of *Atoh1* and *Myo7A* was detected as described above. Expression of tdTOMATO was visualized directly. Labeling was imaged on a Zeiss LSM710.

For western blot analysis, expression constructs for a human *ATOH1*^*WT*^ or *ATOH1*^*c.1030delC*^ were generated in *pCdT* as described above, with 3xFLAG and 3xHA tags were added at the N-terminus of the *ATOH1* sequence. HEK293T cells were transfected with either *pCdT 3xFLAG3xHA-ATOH1*^*WT*^-*IRES-tdTom* or *pCdT 3xFLAG3xHA-ATOH1*^*c.1030delC*^--*IRES-tdTom* using Xfect Transfection Reagent (Clontech). At day 3 after transfection, cells were treated with 1 mM cycloheximide (Sigma) to inhibit new protein synthesis and harvested in cold PBS after 0, 1, 2, 3, and 5 hrs. RIPA buffer supplemented with protease inhibitor cocktail cOmplete Mini (Roche) and phosphatase inhibitor cocktail PhosSTOP (Roche) was used to lyse the cells. After lysis and clarification, the supernatant was divided into two portions – one for BCA protein assay (Thermol) to quantify the total amount of protein, and the other for gel separation using NuPAGE 4-12% SDS-Bis-Tris gel (Invitrogen). After gel separation, the protein was transferred onto a cellulose membrane using an iBlot apparatus (Invitrogen) following the manufacturer’s suggested protocol. Western blotting was carried out using the iBind Flex apparatus (ThermoFisher). Antibodies against the HA tag (H7; Sigma) or ACTIN (Sigma) were used at a 1:5,000-1:10,000 dilution. Binding was detected using an HRP0-conjugated anti-mouse IgG antibody (Jackson Immunoresearch Lab) at a dilution of of 1:5,000-1:10,000. For ECL visualization, Clarity Western ECL substrate A and B solutions (Bio-Rad) were used, and DBio Blue sensitive X-ray film (Vita Scientific) was used for image capture. After X-ray film exposure and development, the bands on the film were scanned at highest resolution (600×600 dpi) on a desktop scanner. ImageJ (NIH) was used to measure the intensity of the bands. A similar area with no band was used as a background control. The reading of ACTIN bands was further used for standardization of the ATOH1 band. For the final presentation, the reading was normalized against the reading at 0 hrs. Two replicates were performed.

#### MITF analysis

For the transactivation assay, HEK293T cells (2×10^4^) were grown for 24 hours in a 96-well plate in 100 μL DMEM + 10% FBS. Cells were transiently co-transfected with an empty pcDNA3.1 construct or pcDNA3.1 containing mouse wild-type *Mitf*^*WT*^ or the *Mitf*^*c.1190delG*^ mutation, a pGL3 basic luciferase construct containing the tyrosinase promotor sequence and a CMV-pRL Renilla control vector. 24 hours post transfection the Dual-Glo Luciferase Assay (Promega) was performed according to the protocol with a Modulus II microplate reader (0.5 sec integration time). Luciferase signals were normalized to corresponding Renilla signals and results expressed as fold change over empty vector. Three biological replicates were performed. For immunoblotting, the cells were lysed and sampled 24 hours post transfection in sample buffer, boiled for 5 min and 20 μL of each sample were analyzed on 8% SDS-polyacrylamide gels that were transferred to nitrocellulose membranes. Membranes were incubated overnight with antibodies for MITF (C5, Thermo Scientific) and β-actin (13E5, Cell Signaling) and then secondary antibodies (anti-rabbit IgG and anti-mouse IgG, Cell Signaling). Results were visualized using the Odyssey Infrared Imaging System.

---

**Figure S1.**
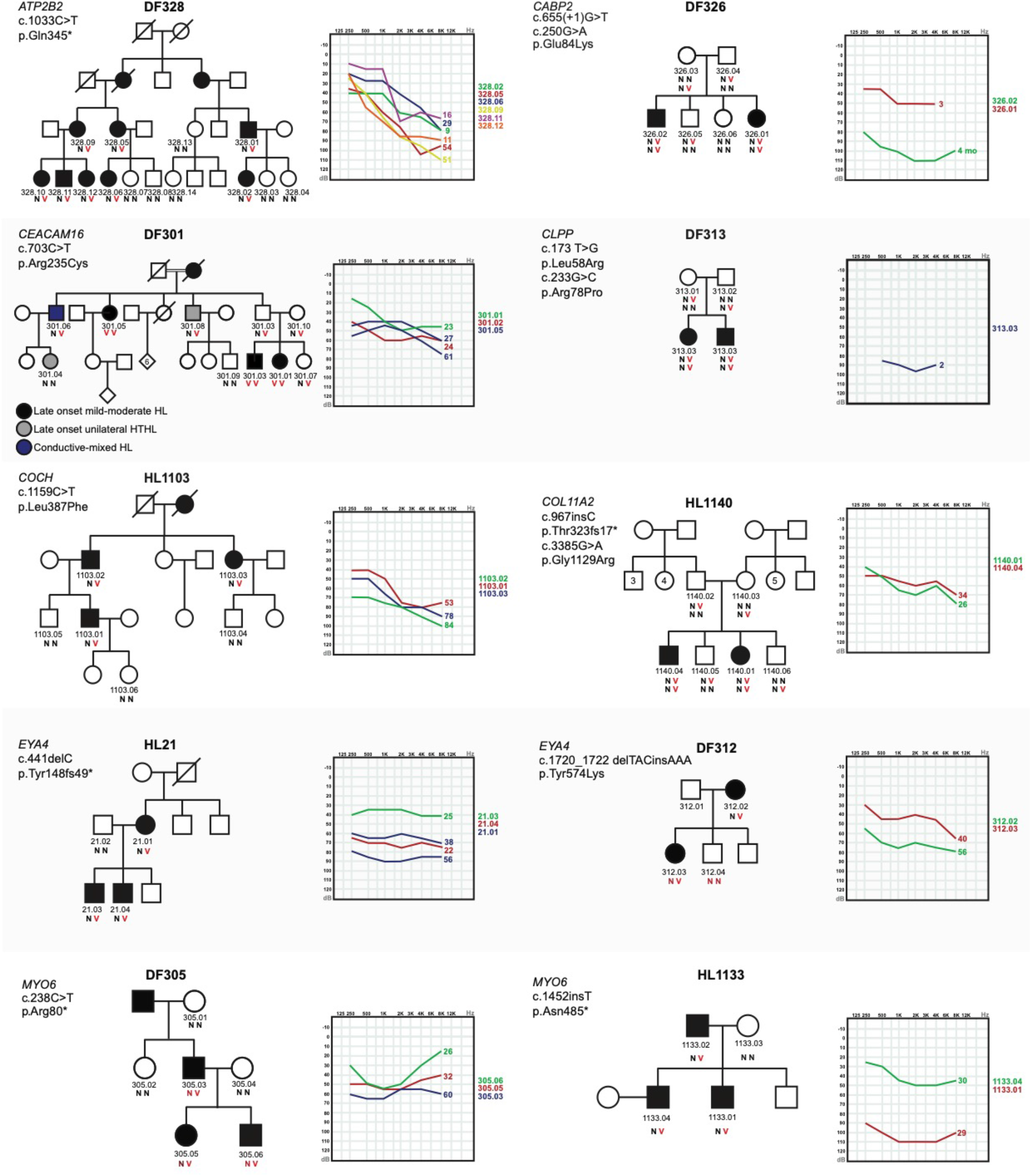
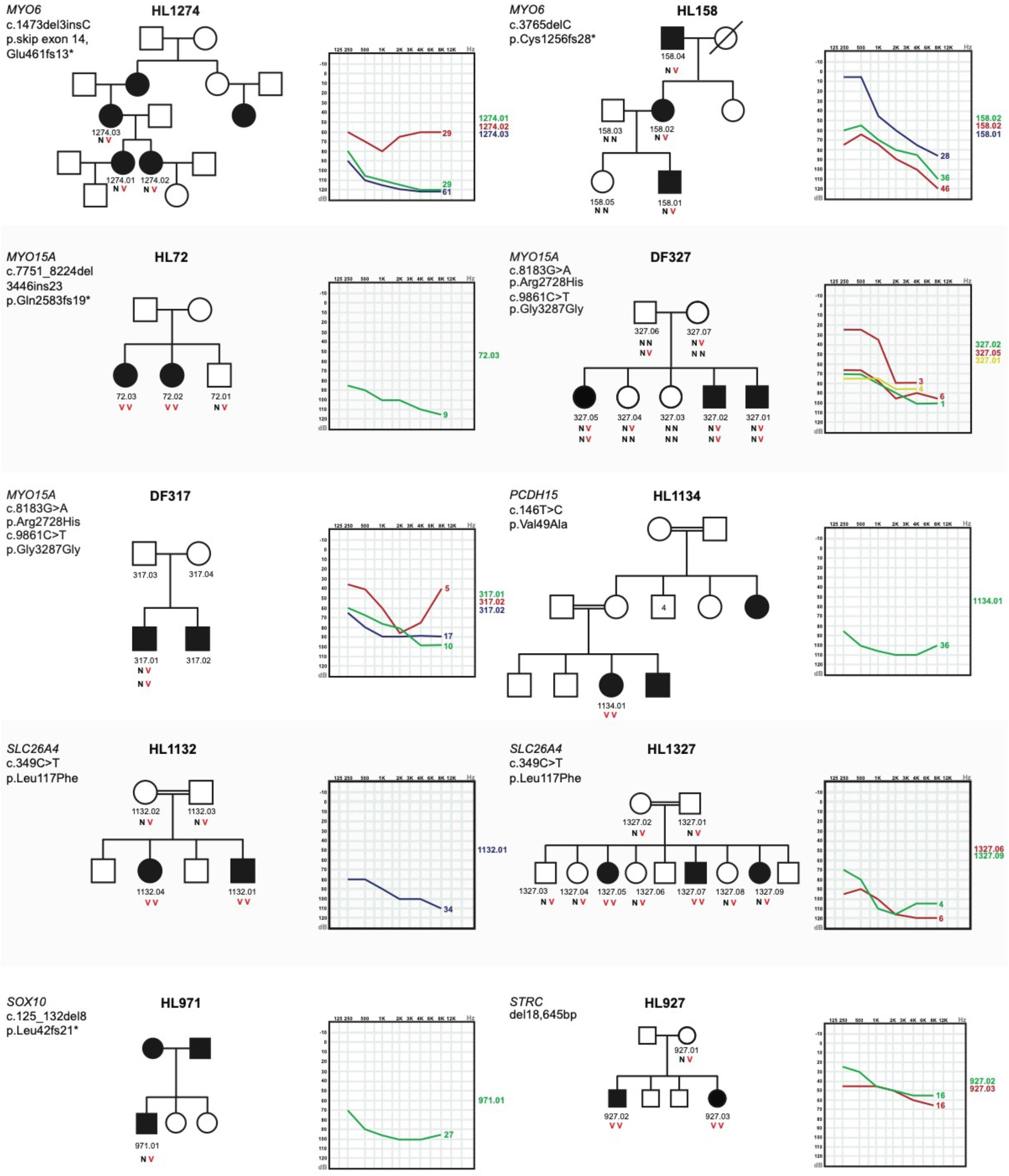
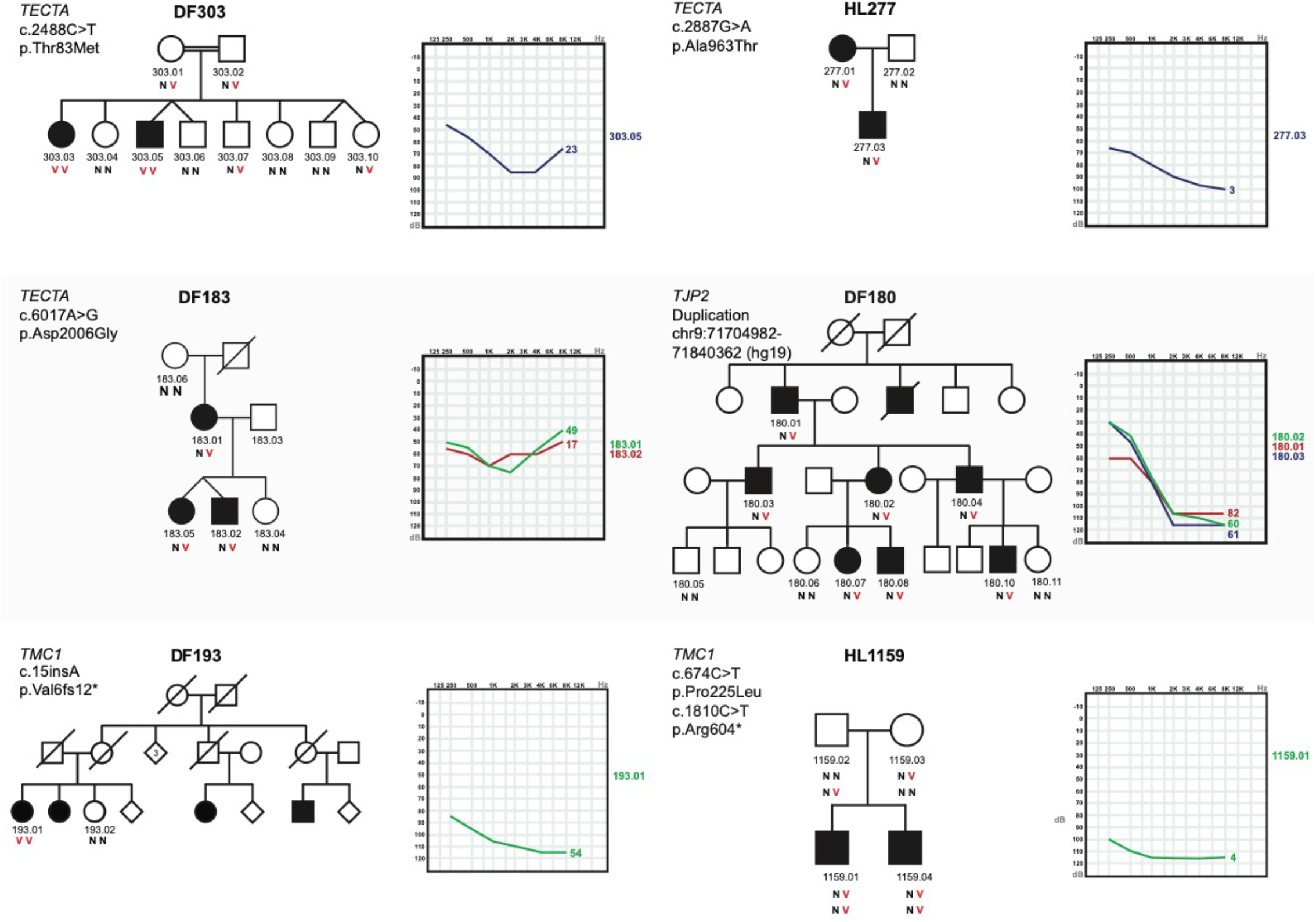
Pedigrees, audiograms and genotypes of families with hearing loss solved by HEar-Seq. Age at the time of testing is noted on audiograms. N = wild type; V = variant.

